# Modeling mesoscopic cortical dynamics using a mean-field model of conductance-based networks of adaptive exponential integrate-and-fire neurons

**DOI:** 10.1101/168385

**Authors:** Yann Zerlaut∗, Sandrine Chemla, Frederic Chavane, Alain Destexhe∗

**Affiliations:** Unité de Neurosciences, Information et Complexité (UNIC). Centre National de la Recherche Scientifique (CNRS), Gif sur Yvette, France.; Neural Coding laboratory. Istituto Italiano di Tecnologia. Rovereto, Italy.; Centre de Recherche Cerveau et Cognition, UMR 5549 CNRS & Université Paul Sabatier Toulouse III, Place du Docteur Baylac, 31059 Toulouse, France.; Institut de Neurosciences de la Timone (INT), UMR 7289 CNRS & Aix-Marseille Université, 27 Bd Jean Moulin, 13385 Marseille cedex 05, France.; The European Institute for Theoretical Neuroscience (EITN), Paris, France.

## Abstract

Voltage-sensitive dye imaging (VSDi) has revealed fundamental properties of neocortical processing at macroscopic scales. Since for each pixel VSDi signals report the average membrane potential over hundreds of neurons, it seems natural to use a mean-field formalism to model such signals. Here, we present a mean-field model of networks of Adaptive Exponential (AdEx) integrate-and-fire neurons, with conductance-based synaptic interactions. We study here a network of regular-spiking (RS) excitatory neurons and fast-spiking (FS) inhibitory neurons. We use a Master Equation formalism, together with a semi-analytic approach to the transfer function of AdEx neurons to describe the average dynamics of the coupled populations. We compare the predictions of this mean-field model to simulated networks of RS-FS cells, first at the level of the spontaneous activity of the network, which is well predicted by the analytical description. Second, we investigate the response of the network to time-varying external input, and show that the mean-field model predicts the response time course of the population. Finally, to model VSDi signals, we consider a one-dimensional ring model made of interconnected RS-FS mean-field units. We found that this model can reproduce the spatio-temporal patterns seen in VSDi of awake monkey visual cortex as a response to local and transient visual stimuli. Conversely, we show that the model allows one to infer physiological parameters from the experimentally-recorded spatio-temporal patterns.

## 1 Introduction

Recent advances in imaging technique, in particular voltage-sensitive dye imaging (VSDi), have revealed fundamental properties of neocortical processing (Arieli et al., 1996; Contreras and Llinas, 2001; Jancke et al., 2004; Ferezou et al., 2006; Chen et al., 2006; Civillico and Contreras, 2012; Muller et al., 2014; Gilad and Slovin, 2015): subthreshold responses to sensory inputs are locally homogeneous in primary sensory areas, depolarizations tend to spread across spatially neighboring regions and responses to sensory stimuli are strongly affected by the level of ongoing activity. It also appears as a great tool to unveil how the spatio-temporal dynamics in the neocortex shape canonical cortical operations such as normalization (Reynaud et al., 2012).

On the other hand, the literature lacks, to the best of our knowledge, theoretical models that provides a detailed account of those phenomena with a clear relation between the biophysical source of the VSDi signal and network dynamics at that spatial scale (i.e. at the millimeters or centimeters scale). Detailed model of a neocortical column (i.e. ~0.5mm^2^ scale) have been recently proposed, see Chemla and Chavane (2010); Chemla and Chavane (2016) for the link with the VSDi signal or more generally Markram et al. (2015), but their computational cost impedes the generalization to higher spatial scale. The aim of the present communication is therefore to design a theoretical model of neocortical dynamics with the following properties: 1) it should have a correlate in terms of single-cell dynamics (in particular membrane potential dynamics), so that the model can directly generate predictions for the signal imaged by the VSDi technique (Berger et al., 2007) and 2) it should describe both the temporal and spatial scale of optical imaging.

More specifically, as we intend to describe responses to salient sensory stimuli, our study focuses on network dynamics in *activated* cortical states(Tan et al., 2014). The desired model should therefore describe neocortical computation in the asynchronous regime, where cortical activity is characterized by irregular firing and strong subthreshold fluctuations at the neuronal level (Steriade et al., 2001; Destexhe et al., 2003). The strategy behind the present model is to take advantage of the *mean-field* descriptions of network dynamics in this regime. Via self-consistent approaches, those descriptions allow to capture the dynamical properties of population activity in recurrent networks (Amit and Brunel, 1997; Brunel and Hakim, 1999; Brunel, 2000; Latham et al., 2000; El Boustani and Destexhe, 2009). The present model thus relies on the following scheme: 1) we consider the randomly connected network of 10000 neurons as a unit to describe few cortical columns and 2) we embedded the analytical description of this cortical column model into a ring geometry with physiological connectivity profiles to model spatio-temporal integration on the neocortical sheet.

We first compare the analytical prediction of the model with numerical simulations in order to evaluate the accuracy and/or weaknesses of our specific analytical description (adapted from (El Boustani and Destexhe, 2009)). We next investigate the integrative properties of the model, i.e. the relation between the network response and the properties of the input. Finally, based on this mean-field approach, we construct a spatio-temporal model for the dynamics of superficial layers in neocortex by arranging mean-field units according to a one-dimentional ring structure. We then compare this model’s response to afferent inputs with VSDi recordings in the primary visual cortex (V1) of awake monkey in response to a visual stimulus.

## 2 Material and Methods

Here, we describe the equations and parameters used for the neuronal, synaptic and network modeling. We present our *heuristic* treatment of the neuronal *transfer functions*: the quantity that accounts for the cellular computation in *mean-field* models of population activity. Then, we present the specific markovian model of population activity used in this study and we construct a spatio-temporal model of neocortical integration by embedding this description into a one-dimensional ring model.

### 2.1 Single neuron models

The neuronal model used in this study is the adaptative exponential and fire (AdEx) model (Brette and Gerstner, 2005). The equation for the membrane potential and the adaptation current therefore reads:

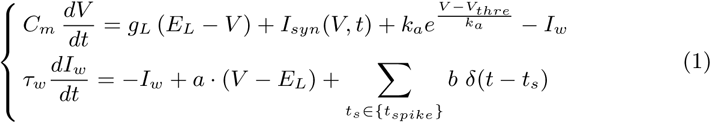

where *I*_*syn*_(*V, t*) is the current emulating synaptic activity that will create the fluctuations, *I*_*w*_ accounts for the phenomena of spike-frequency adaptation as well as subthreshold adaptation(McCormick et al., 1985). The spiking mechanism is the following: a spike is triggered at t_s_ ϵ {t_spike_} when *V* (*t*) reaches *V*_*thre*_ +5 *k*_*a*_. Afterwards, the adaptation variable *I*_*w*_ is incremented by *b* and the membrane potential is then clamped at *E*_*L*_ for a duration *τ* _refrac_=5ms. We consider two versions of this model: a regular spiking neuron for the excitatory cells and a fast spiking neuron for the inhibitory cells (see Fig. 2). The parameters of those two models can be found on Table 1.

**Table 1:**
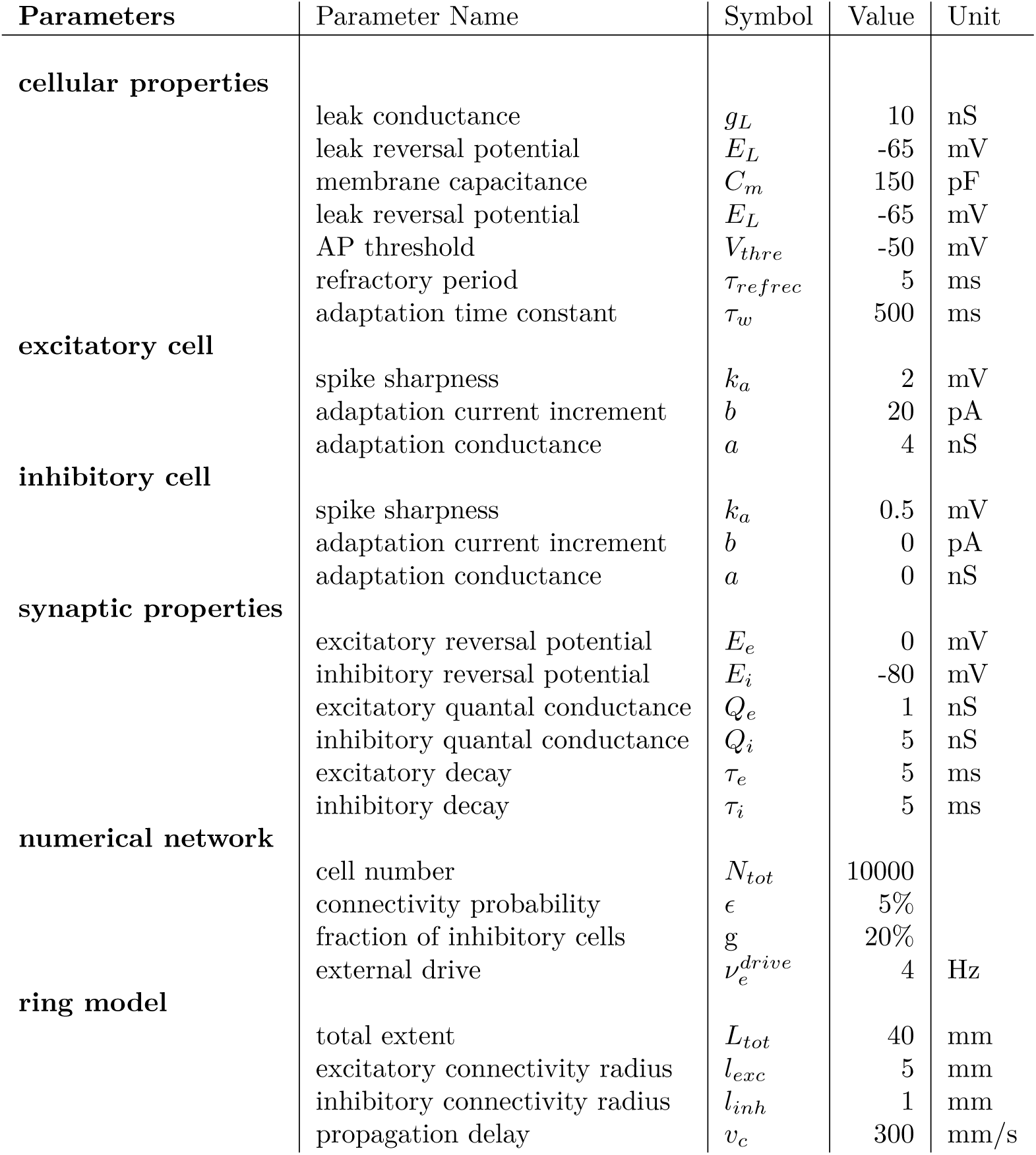
Model parameters.

| Parameters | Parameter Name | Symbol | Value | Unit |
| --- | --- | --- | --- | --- |
| cellular properties | leak conductance | $g_L$ | 10 | nS |
| | leak reversal potential | $E_L$ | -65 | mV |
| | membrane capacitance | $C_m$ | 150 | pF |
| | leak reversal potential | $E_L$ | -65 | mV |
| | AP threshold | $V_{thre}$ | -50 | mV |
| | refractory period | $\tau_{refrec}$ | 5 | ms |
| | adaptation time constant | $\tau_w$ | 500 | ms |
| excitatory cell | spike sharpness | $k_a$ | 2 | mV |
| | adaptation current increment | $b$ | 20 | pA |
| | adaptation conductance | $a$ | 4 | nS |
| inhibitory cell | spike sharpness | $k_a$ | 0.5 | mV |
| | adaptation current increment | $b$ | 0 | pA |
| | adaptation conductance | $a$ | 0 | nS |
| synaptic properties | excitatory reversal potential | $E_e$ | 0 | mV |
| | inhibitory reversal potential | $E_i$ | -80 | mV |
| | excitatory quantal conductance | $Q_e$ | 1 | nS |
| | inhibitory quantal conductance | $Q_i$ | 5 | nS |
| | excitatory decay | $\tau_e$ | 5 | ms |
| | inhibitory decay | $\tau_i$ | 5 | ms |
| numerical network | cell number | $N_{tot}$ | 10000 | |
| | connectivity probability | $\epsilon$ | 5% | |
| | fraction of inhibitory cells | $g$ | 20% | |
| | external drive | $\nu_e^{drive}$ | 4 | Hz |
| ring model | total extent | $L_{tot}$ | 40 | mm |
| | excitatory connectivity radius | $l_{exc}$ | 5 | mm |
| | inhibitory connectivity radius | $l_{inh}$ | 1 | mm |
| | propagation delay | $v_c$ | 300 | mm/s |

### 2.2 Synaptic model

The time- and voltage-dependent current that stimulate the neuron is made of the sum of excitatory and inhibitory currents (indexed by *s* ϵ {*e, i*} and having a reversal potential *E*_*s*_):

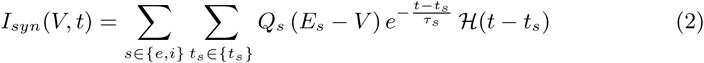

where 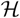 is the Heaviside function.

This synaptic model is referred to as the *conductance-based exponential* synapse. The set of events {*t*_*e*_} and {*t*_*i*_} are the set of excitatory and inhibitory events arriving to the neuron. In numerical simulations of single neurons (performed to determine the *transfer function* 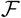 of either excitatory or inhibitory neurons), it will be generated by stationary Poisson processes. On the other hand, in numerical simulations of network dynamics it will correspond to the set of spike times of the neurons connecting to the target neurons, both via recurrent and feedforward connectivity.

### 2.3 Numerical network model

All simulations of numerical network were performed with the brian2 simulator (Goodman and Brette, 2009), see http://brian2.readthedocs.org. For all simulations, the network was composed of *N*_*tot*_=10000 neurons, separated in two populations, one excitatory and one inhibitory with a ratio of g=20% inhibitory cells. Those two local populations were randomly connected (internally and mutually) with a connectivity probability *ε*=5%.

Because this network did not display self-sustained activity (in contrast to Vogels and Abbott (2005)), an excitatory population exerted an *external drive* to bring the network out of the quiescent state. This population targeted both the excitatory and inhibitory neurons. Note that the firing rate of this population was linearly increased to avoid a too strong initial synchronization (see Fig. 3). Finally, when studying responses to external inputs, an excitatory population of time varying firing rate was added to evoke activity transients in the population dynamics. This last stimulation targeted only the excitatory population. The number of neurons in those two excitatory populations was taken as identical to the number of excitatory neurons (i.e. (1 − *g*) *N*_*tot*_) and created synapses onto the recurrent network with the same probability *ϵ*. After temporal discretization, the firing rates of those afferent populations were converted into spikes by using the properties of a Poisson process (i.e. eliciting a spike at *t* with a probability *ν*(*t*) *dt*). All simulations were performed with a time-step *dt*=0.1ms.

**Figure 1:**
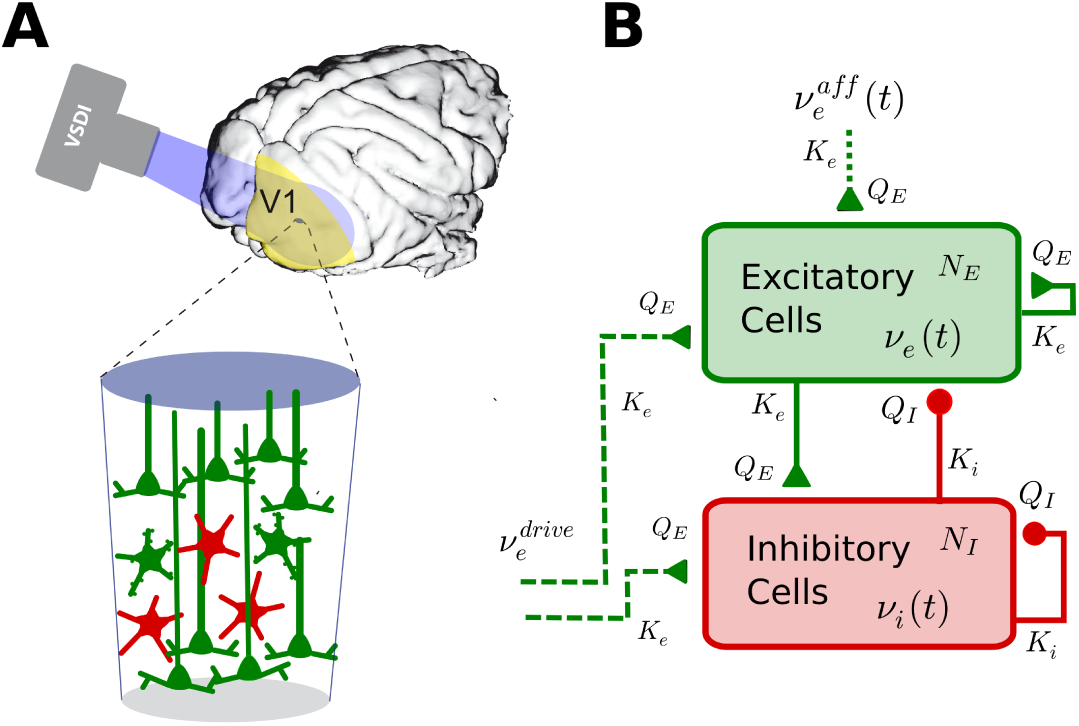
Modeling local cortical dynamics. **(A)** The complex cellular assembly corresponding to a single pixel in VSD imaging is reduced to a local excitatory-inhibitory network. **(B)** Schematic of the local network architecture. The network is made of *N*_*e*_ = (1 − *g*) *N*_*tot*_ excitatory and *N*_*i*_ = *g N*_*tot*_ inhibitory neurons. All excitatory connections (afferent and recurrent) onto a neuron corresponds to *K*_*e*_ = *ε* (1 − *g*) *N*_*tot*_ synapses of weight *Q*_*e*_. All inhibitory connections onto a neuron corresponds to *K*_*i*_ = *ε g N*_*tot*_ synapses of weight *Q*_*i*_.

### 2.4 Estimating the transfer functions of single neurons

The transfer function 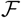 of a single neuron is defined here as the function that maps the value of the stationary excitatory and inhibitory presynaptic release frequencies to the output stationary firing rate response, i.e. 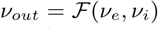. Note the stationary hypothesis in the definition of the transfer function (see discussion in main text).

Because an analytical solution (of this function 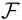) for the single neuron models considered in our study is a very challenging mathematical problem, we adopted a semi-analytical approach. We performed numerical simulations of single cell dynamics at various excitatory and inhibitory presynaptic frequencies (*ν*_*e*_ and *ν*_*i*_ respectively) (see the output in Fig. 2) on which we fitted the coefficients of an analytical template to capture the single cell model’s response.

**Figure 2:**
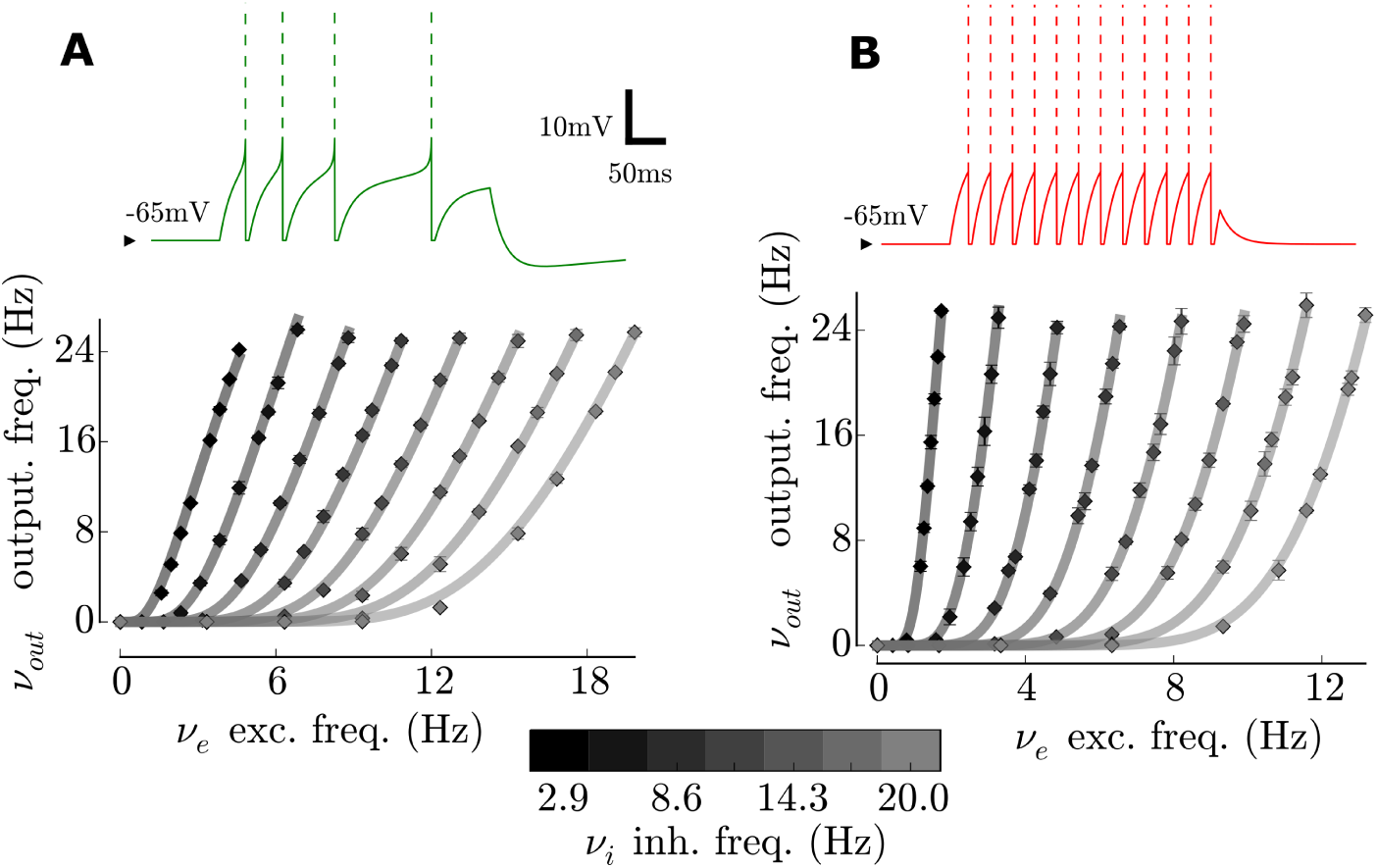
Single cell models of the excitatory and inhibitory populations. Top: response to a current step of 200pA lasting 300ms. Bottom: *transfer function* of the single cell, i.e. output firing rate as a function of the excitatory (x-axis) and inhibitory (color-coded) presynaptic release frequencies. Note that the range of the excitatory and frequencies assumes numbers of synapses *K*_*e*_ = *ϵ* (1 – *g*) *N*_*tot*_ and *K*_*i*_ = *ϵ g N*_*tot*_ for the excitation and inhibition respectively). **(A)** Excitatory cells. Note the presence of spike-frequency adaptation and subthreshold adaptation. **(B)** Inhibitory cells. Note the very narrow spike initiation dynamics. Also, note the steepest relation to excitation (with respect to the excitatory cell) at various inhibitory levels as a result of the increased excitability of the inhibitory cell (with respect to the excitatory cell).

The procedure relied on fitting a *phenomenological threshold* 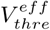 that accounts for the single neuron non-linearities (spiking and reset mechanism, adaptation mechanisms) on top of the subthreshold integration effects (Zerlaut et al., 2016). This phenomenological threshold is then plugged-in into the following formula (analogous to Amit and Brunel (1997)) to become our firing response estimate:

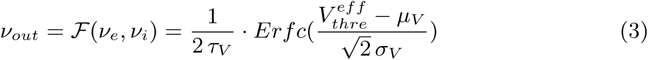

Where (*µ*_*V*_, *σ*_*V*_, *τ*_*V*_) are the mean, standard deviation and autocorrelation time constant of the membrane potential fluctuations. How to calculate those quantities as a response to a stationary stimulation is the focus of the next section.

The expression for the phenomenological threshold was the following:

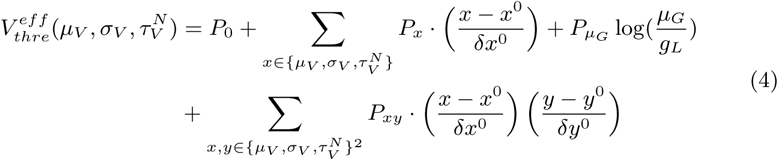

We took a second order polynomial in the three dimensional space (*µ*_*V*_, *σ*_*V*_, *τ*_*V*_) combined with a term capturing the effect of total conductance on the effective threshold (Platkiewicz and Brette, 2010). The normalization factors 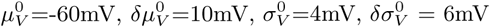 *τ*_*V*_ =10ms and *δτ*_*V*_ = 20ms arbitrarily delimits the *fluctuation-driven* regime (a mean value *x* and an extent *δx*, 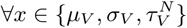). They render the fitting of the phenomenological threshold easier, as they insure that the coefficients take similar values. It is kept constant all along the study. The phenomenological threshold was taken as a second order polynomial and not as a linear threshold, for two reasons: 1) unlike in an experimental study (Zerlaut et al., 2016), we are not limited by the number of sampling points, the number of fitted coefficients can thus be higher as the probability of overfitting becomes negligible 2) it gives more flexibility to the template, indeed the linear threshold was found a good approximation in the *fluctuation-driven* regime (Zerlaut et al., 2016), i.e. when the diffusion approximation holds. However, for low values of the presynaptic frequencies, we can be far from this approximation, the additional coefficients are used to capture the firing response in those domains. Those coefficients are listed on Table 2 for the two cell types (RS & FS).

**Table 2:**
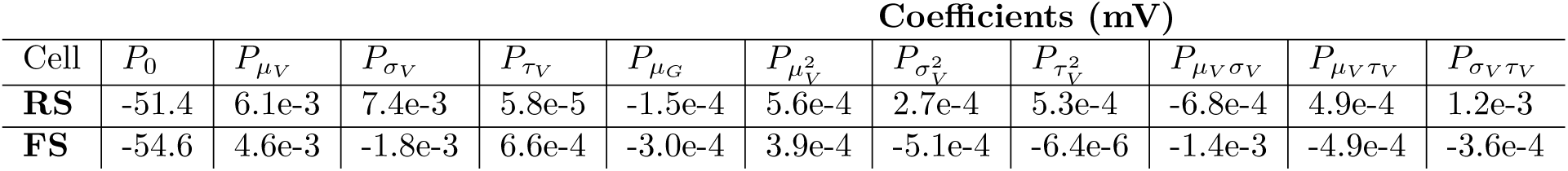
Fitted coefficients of the transfer functions (see Fig. 2).

| Cell | Coefficients (mV) |  |  |  |  |  |  |  |  |  |  |
| --- | --- | --- | --- | --- | --- | --- | --- | --- | --- | --- | --- |
| | $P_0$ | $P_{\mu_V}$ | $P_{\sigma_V}$ | $P_{\tau_V}$ | $P_{\mu_G}$ | $P_{\mu_V^2}$ | $P_{\sigma_V^2}$ | $P_{\tau_V^2}$ | $P_{\mu_V \sigma_V}$ | $P_{\mu_V \tau_V}$ | $P_{\sigma_V \tau_V}$ |
| **RS** | -51.4 | 6.1e-3 | 7.4e-3 | 5.8e-5 | -1.5e-4 | 5.6e-4 | 2.7e-4 | 5.3e-4 | -6.8e-4 | 4.9e-4 | 1.2e-3 |
| **FS** | -54.6 | 4.6e-3 | -1.8e-3 | 6.6e-4 | -3.0e-4 | 3.9e-4 | -5.1e-4 | -6.4e-6 | -1.4e-3 | -4.9e-4 | -3.6e-4 |

The fitting procedure was identical to Zerlaut et al. (2016), it consisted first in a linear regression in the phenomenological threshold space of Equation 4, followed by a non-linear optimization of Equation 3 on the firing rate response. Both fitting were performed with the `leastsq` method in the `optimize` package of `SciPy`.

### 2.5 Calculus of the subthreshold membrane potential fluctuations

Here, we detail the analytical calculus that translate the input to the neuron into the properties of the membrane potential fluctuations. The input is made of two Poisson shotnoise: one excitatory and one inhibitory that are both convoluted with an exponential waveform to produce the synaptic conductances time courses.

#### 2.5.1 Conductances fluctuations

From Campbell’s theorem (Papoulis, 1991), we first get the mean (*µ*_*Ge*_, *µ*_*Gi*_) and standard deviation (*σ*_*Ge*_, *σ*_*Gi*_) of the excitatory and inhibitory conductance:

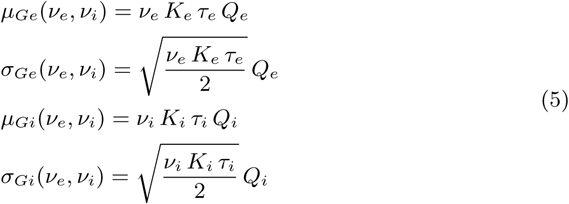

The mean conductances will control the input conductance of the neuron *µ*_*G*_ and therefore its effective membrane time constant *τ*_*m*_:

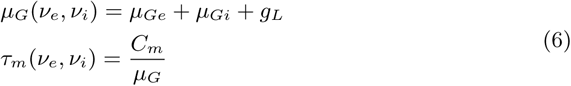

#### 2.5.2 Mean membrane potential

Following Kuhn et al. (2004), the mean membrane potential is obtained by taking the stationary solution to static conductances given by the mean synaptic bombardment (for the passive version of Equation 1, i.e. removing the adaptation and spiking mechanisms). We obtain:

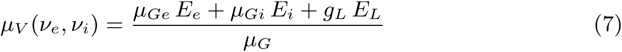

We will now approximate the driving force *E*_*s*_ − *V* (*t*) of synaptic events by the level resulting from the mean conductance bombardment: *E*_*s*_ − *µ*_*V*_. This will enable an analytical solution for the standard deviation *σ*_*V*_ and the autocorrelation time *σ*_*V*_ of the fluctuations.

#### 2.5.3 Power spectrum of the membrane potential fluctuations

Obtaining *σ*_*V*_ and *τ*_*V*_ is achieved by computing the power spectrum density of the fluctuations. In the case of Poisson processes, the power spectrum density of the fluctuations resulting from the sum of events *PSP*_*s*_(*t*) at frequency *K*_*s*_*ν*_*s*_ can be obtained from shotnoise theory (Daley and Vere-Jones, 2007):

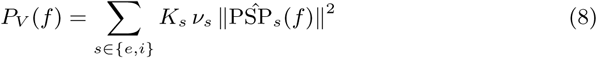

where 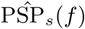 is the Fourier transform of the time-varying function PSP(*t*). Note that the relations presented in this paper rely on the following convention for the Fourier transform: 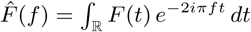.

After fixing the driving force to *E*_*s*_ − *μ*_*V*_, the equation for a post-synaptic membrane potential event *s* around *μ*_*V*_ is:

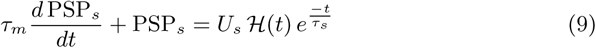

where 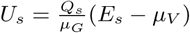 and 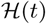 is the Heaviside function.

Its solution is:

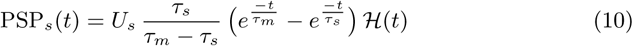

We take the Fourier transform:

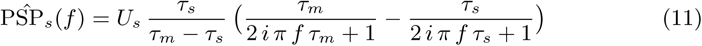

We will need the value of the square modulus at *f* = 0:

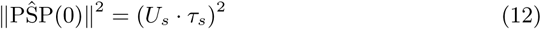

As well as the integral of the square modulus:

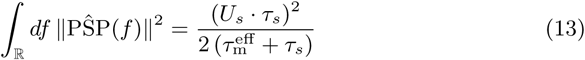

#### 2.5.4 Standard deviation of the fluctuations

The standard deviation follows:

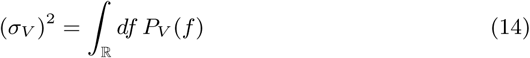

Using Equation 13, we find the final expression for *σ*_*V*_:

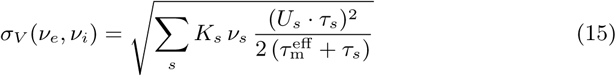

#### 2.5.5 Autocorrelation-time of the fluctuations

We defined the global autocorrelation time as (Zerlaut et al., 2016):

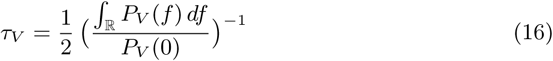

Using Equations 13 and 12, we find the final expression for *τ*_*V*_:

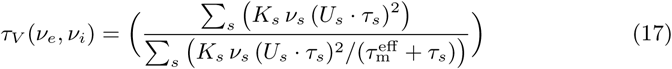

Therefore the set of Equations 7, 15 and 17 translates the presynaptic frequencies into membrane fluctuations properties *µ*_*V*_, *σ*_*V*_, *τ*_*V*_.

The previous methodological section allowed to translate the fluctuations properties *µ*_*V*_, *σ*_*V*_, *τ*_*V*_ into a spiking probability thanks to a minimization procedure. The combination of the present analytical calculus and the previous fitting procedure (on numerical simulations data) constitute our semi-analytical approach to determine the transfer function of a single cell model: 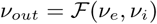.

### 2.6 Master equation for local population dynamics

As we now benefit from an analytical description of the cellular transfer function, we can now turn to the theoretical analysis of asynchronous dynamics in sparsely connected random networks (Amit and Brunel, 1997; Brunel, 2000; Renart et al., 2004).

Because we will investigate relatively slow dynamics (*τ*>25-50ms) (and because of the stationary formulation of our transfer function), we will use the Markovian description developed in El Boustani and Destexhe (2009), it describes network activity at a time scale *T*, for which the network dynamics should be Markovian. The choice of the time-scale *T* is quite crucial in this formalism, it should be large enough so that activity can be considered as memoryless (e.g. it can not be much smaller than the refractory period, that would introduce memory effects) and small enough so that each neuron can fire statistically less than once per time interval *T*. Following El Boustani and Destexhe (2009), we will arbitrarily take *T*=5ms all along the study as it offers a good compromise between those two constraints.

The formalism describes the first and second moments of the population activity for each populations. We consider here two populations: one excitatory and one inhibitory, the formalism thus describes the evolution of five quantities: the two means *ν*_*e*_(*t*) and *ν*_*i*_(*t*) of the excitatory and inhibitory population activity respectively (the instantaneous population firing rate, i.e. after binning in bins of *T*=5ms, see discussion in El Boustani and Destexhe (2009)), the two variances *c*_*ee*_(*t*) and *c*_*ii*_(*t*) of the the excitatory and inhibitory population activity respectively and the covariance *c*_*ei*_(*t*) between the excitatory and inhibitory population activities. The set of differential equations followed by those quantities reads (El Boustani and Destexhe, 2009):

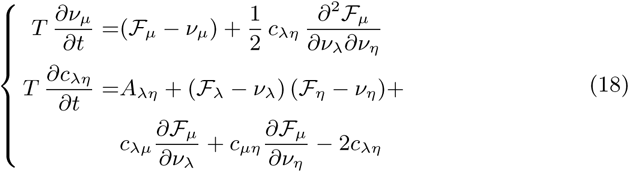

with:

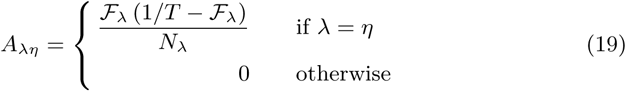

Note that, for the concision of the expressions, we used Einstein’s index summation convention: if an index is repeated in a product, a summation over the whole range of value is implied (e.g. we sum over *λ* ϵ {*e, i*} in the first equation, consequently, *λ* does not appear in the left side of the equation). Also the dependency of the firing rate response to the excitatory and inhibitory activities has been omitted: yielding 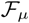 instead of 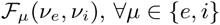. We will also use the reduction to first order of this system (for the ring model). This yields:

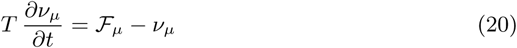

### 2.7 Afferent stimulation

The afferent input was represented by the following piecewise double Gaussian waveform:

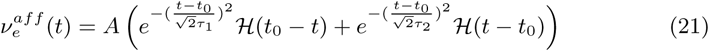

In this afferent input, we can independently control: 1) the maximum amplitude *A* of the stimulation, its rising time constant *τ*_1_ and its decay time constant *τ*_2_.

### 2.8 Ring model

To model VSDi experiments (See Fig. 1A), we embed the mean-field description of population dynamics in a ring geometry to model spatio-temporal integration on the neocortical sheet. The ring geometry corresponds to a one dimensional spatial description with an invariance by translation, i.e. for all quantities *f*, *f*(*x*) = *f*(*x* + *L*), also termed one dimensional periodic boundary conditions, where *L* is the length of the ring model. For simplicity, we consider here only the first moments of the second-order description: i.e. the means of the excitatory and inhibitory population activities: *ν*_*e*_(*t*) and *ν*_*i*_(*t*) respectively.

We use a Gaussian connectivity profile (see Fig. 7) to define the connectivity across cortical columns (i.e. local networks described in the previous section):

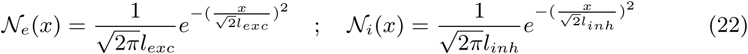

where *l*_*exc*_ and *l*_*inh*_ are the excitatory and inhibitory extent of the connectivity profiles respectively.

We also introduce the effect of a finite axonal conduction speed *v*_*c*_, this will introduce delays for the propagation of activity across cortical columns: for a network at a distance *x*, the afferent activity will arrive delayed by *x*/*v*_*c*_.

Finally, the equations that govern the activity in space and time are given by:

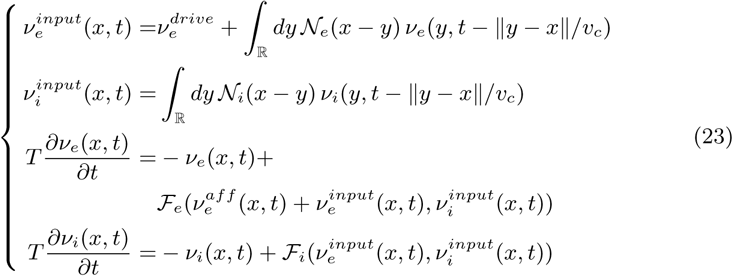

where 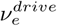 is the external drive and 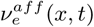 (*x, t*) is the afferent (thalamic) stimulation.

The local correlate in terms of mean membrane potential *µ*_*V*_(*x, t*) is given by Equation 7. Because VSDi provides a variation with respect to the fluorescence baseline (Berger et al., 2007) (i.e. the relative membrane potential deflection of a population with respect to mean the membrane potential at the level of spontaneous network activity), we also present the variations of a normalized membrane potential quantity:

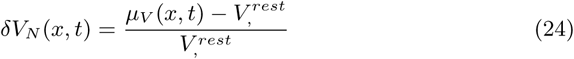

where *V*^*rest*^ is the mean membrane potential during spontaneous activity in the model (see Fig. 3).

### 2.9 Visually-evoked patterns of cortical activity in awake monkey recorded through Voltage-Sensitive Dye imaging

The experimental procedure was described elsewhere (Reynaud et al., 2012; Muller et al., 2014), briefly summarized below

Surgical preparations: Experiments were conducted on two male rhesus monkeys (macaca mulatta, aged 14 and 11 years old respectively for monkey WA and monkey BR). The monkeys were chronically implanted with a head-holder and a recording chamber located above the cortical areas V1 and V2 of the right hemisphere. The dura was surgically removed over a surface corresponding to the recording aperture (18 mm diameter) and a silicon-made artificial dura was inserted under aseptic conditions (Arieli et al., 2002). Before each recording session, the cortex was stained with the voltage-sensitive dye RH-1691 (Optical Imaging), prepared in artificial cerebrospinal fluid (aCSF) at a concentration of 0.2 mg/ml and filtered through a 0.2 *µ*m filter. During the recordings, the cortex was illuminated at 630 nm to excite the dye for 700 ms. Experimental protocols have been approved by the Marseille Ethical Committee in Neuroscience (approval A10/01/13, official national registration French Ministry of Research). All procedures complied with the French and European regulations for animal research, as well as the guidelines from the Society for Neuroscience.

VSD imaging protocol. Experimental controls and online eye position monitoring were performed by a PC running the REX software (NEI-NIH) with the QNX operating system. Eye position was monitored with an ISCAN at 1kHz (ETL-200 Eye Tracking System). Optical signals were recorded with a Dalstar camera (512 x 512 pixels, 110 Hz frame rate) driven by the Imager 3001 system (Optical Imaging). Both online behavioural control and image acquisition were heartbeat regulated. Heartbeat was detected with a pulse oximeter (Nonin 8600V). The visual stimuli were computed online using VSG2/5 libraries and were displayed on a 22 inch CRT monitor at a resolution of 1,024 x 768 pixels. Refresh rate was set to 100 Hz. Viewing distance was 57 cm. Luminance values were linearized by means of a look-up table.

Visual stimuli. During a single trial, the monkey had to fixate on a central red dot for 1–2 s. The animal’s gaze was constrained in a window of 2°x 2°. Stimuli were presented during fixation, and a reward (water or applesauce drop) was given after the trial if the monkey maintained fixation during the acquisition period. Each trial ran for 700 ms: 100-200ms delay, 20–100ms stimulation period, 300-580ms post-stimulus period. Stimuli were local Gaussian blobs with a s.d. of 0.5° in space. Stimuli were presented at 0.5° on the left of the vertical meridian and 3° below the horizontal, except where noted. Three different durations were used: 20, 50, 100 ms.

Data analysis. Stacks of images were stored on hard-drives for offline analysis with MATLAB R2014a (MathWorks), using the Optimization, Statistics, and Signal Processing Toolboxes. The evoked response to each stimulus was computed in four successive basic steps. First, the recorded value at each pixel was divided by the average value before stimulus onset (“frames 0 division”) to remove slow stimulus-independent fluctuations in illumination and background fluorescence levels. Second, this value was subsequently subtracted by the value obtained for the blank condition (“blank subtraction”) to eliminate most of the noise due to heartbeat and respiration (Shoham et al., 1999). Third, a linear detrending of the time series was applied to remove residual slow drifts induced by dye bleaching (Chen et al., 2008; Meirovithz et al., 2009).

### 2.10 Optimizing model parameters with respect to single-session VSDi recordings

We describe here the optimization procedure used to estimate a set of model parameters for each of the 12 VSDi recording sessions described above. We optimized the following parameters of the ring model: the speed of lateral propagation *v*_*c*_, the spatial extent of the excitatory connectivity *l*_*exc*_, the spatial extent of the inhibitory connectivity *l*_*inh*_, the spatial extent of the stimulus *l*_*stim*_, the onset time constant of the stimulus *τ*_1_ and the decay time constant of the stimulus *tau*_2_. The three first parameters (*v*_*c*_, *l*_*exc*_ and *l*_*inh*_) are spatial parameters of the dynamics (i.e. they enter into Equation 26), whereas the three last parameters (*l*_*stim*_, *τ*_1_ and *τ*_2_) are the spatial and temporal parameters of the afferent input (i.e. they enter into Equations 22 and 23). Parameter scans were performed on a grid of parameters with the following boundaries: *v*_*c*_ ϵ [50, 600]mm/s, *l*_*exc*_ ϵ [1, 7]mm, *l*_*inh*_ ϵ [1, 7]mm, *τ*_1_ ϵ [5, 50]ms,*τ*_2_ ϵ [50, 200]ms. We scanned 5 points per dimension, i.e. we simulated and analyzed 15625 configurations.

For a given session (see examples in Fig. 9B), the optimal parameters of the model were taken as those minimizing the square residual of the difference between spatio-temporal pattern. To obtain the residual of the difference between the model and VSDi data, one needs to have a common sampling of the pattern. We thus subsampled the model data over time to reach the sampling rate of the VSDi data (i.e. *f*_*smpl*_=110Hz). Data were aligned with respect to the local maximum over space and time (*t*_*center*_, *x*_*center*_). The temporal window was taken as a 400ms window, with 100ms before *t*_*center*_ and 300ms after *t*_*center*_. The spatial sampling of the model was reduced to the field of view of the VSDi recording (that varies over session, see the different *x*_*center*_ locations in Fig. 9B). VSDi data were normalized per session with respect to their maximum, resulting in the signal 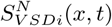 with values below 1. The model data were all normalized to a common factor *δµ*_*V*_ =7% (i.e. implementing a physiological constraint: average evoked depolarizations should reach the 55mV range to match VSDi data normalization), resulting in the signal 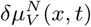. On Fig. 9B, one can see examples of the 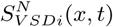 and 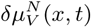 signals corresponding to a single session. Given this normalization rules and the common spatio-temporal sampling, the residual was simply given by:

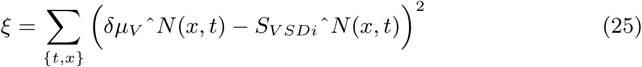

## 3 Results

The results are organized as follows. We construct the analytical model that describes the dynamics of a population of RS and FS cells. We start by describing the semi-analytical workflow that enables the derivation of the cellular transfer function: the core of this population model. Next, we investigate whether the analytical description accurately describe population dynamics by comparing its prediction to numerical simulations. We also investigate the response of the network model subject to an external input. Finally, we build a 1-dimensional ring model made of interconnected RS-FS mean-field units and investigate if this model can reproduce the visually-evoked patterns of activity seen in VSDi of awake monkey visual cortex.

### 3.1 Modeling a local cortical population

We adopt a simplistic description of a local cortical population (see Fig. 1). The complex cellular assembly is reduced to a two population network model: one excitatory and one inhibitory comprising 8000 and 2000 neurons respectively. All neurons within the two population synaptically interconnect randomly to each other with a connectivity probability of 5%. The excitatory and inhibitory cells have the same passive properties. We nonetheless include an asymmetry between the excitatory and inhibitory populations: because the inhibitory population includes Fast-Spiking cells that can exhibit very high firing frequencies (Markram et al., 2004), we set its spiking mechanism sharper (more precisely its sodium activation activation curve is steeper, see Methods) than that of excitatory cells, additionally we add a strong spike-frequency adaptation current in excitatory cells that is absent in inhibitory cells. Those two effects render the inhibitory neurons more excitable (see the different responses to the same current step in Fig. 2). All parameters of the cortical column can be found in Table 1.

### 3.2 A Markovian model to describe population dynamics

We now want to have an analytical description of the collective dynamics of this local network. We adopted the formalism presented in El Boustani and Destexhe (2009). Two reasons motivated this choice: 1) because 10000 neurons is still far from the large network limit, finite-size effects could have a significant impact on the dynamics and 2) because of the relative complexity of the cellular models, an analytic treatment of the type Amit and Brunel (1997) is, to our knowledge, not accessible and would be extremely challenging to derive. The Markovian framework proposed in El Boustani and Destexhe (2009) positively respond to those two constraints: it is a second-order description of population activity that describes fluctuations emerging from finite-size effects and it is applicable to any neuron model as long as its transfer function can be characterized. In a companion study (Zerlaut et al., 2016), we developed a semi-analytical approach to characterize those transfer functions (see next section), we will therefore incorporate this description into the formalism.

Nonetheless, the study of El Boustani and Destexhe (2009) only investigated the ability of the formalism to describe 1) the stationary point of the network activity and 2) in a situation where the neuronal models models had an analytic estimate for the transfer function (current-based integrate-and-fire model). As a prerequisite, investigating whether this description generalizes to transient dynamics and transfer functions estimated with a semi-analytical approach is investigated in the next sections.

### 3.3 Transfer functions of excitatory and inhibitory cells

We briefly describe here the semi-analytical approach used to characterize the transfer function (see details in the Methods). The transfer function 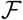 of a single neuron is defined here as the function that maps the value of the stationary excitatory and inhibitory presynaptic release frequencies to the output stationary firing rate response, i.e. 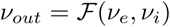. This kind of input-output functions lie at the core of *mean-field* models of population dynamics (reviewed in Renart et al. (2004)) and is consequently the main ingredient of the formalism adopted here (El Boustani and Destexhe, 2009). Note here that the formulation of the transfer function imply a stationary hypothesis: both for the input (stationary Poisson processes) and the output firing (a stationary firing rate). We will study in the following what are the limitations introduced by this stationary hypothesis in the description of the temporal dynamics of network activity.

In a previous communication (Zerlaut et al., 2016), we found that the firing rate response of several models (including the adaptative exponential integrate and fire considered in this study) would be captured by a *fluctuations-dependent* threshold in a simple approximation of the firing probability (see Methods).

The semi-analytical approach thus consisted in making numerical simulations of single-cell dynamics for various presynaptic activity levels (i.e. scanning various *ν*_*e*_, *ν*_*i*_ configurations) and measuring the output firing rate *ν*_*out*_. All those configurations corresponded to analytical estimates of (*µ*_*V*_, *σ*_*V*_, *τ*_*V*_), we then fitted the *fluctuations-dependent* threshold that bring the analytical estimate to the measured firing response. This procedure resulted in the analytical estimates shown in Fig. 2 and compared with the results of numerical simulations.

### 3.4 Accuracy of the description of the spontaneous activity state

We now compare the numerical simulation of the network model (Fig. 3) to the prediction of the Markovian description in terms of stationary dynamics. First, we see that there is a transient period of ~ 400ms resulting from the onset of the external drive (see Fig. 3B-D), we will therefore evaluate stationary properties after discarding the first 500ms of the simulation. After this initial transient, the population activities (*ν*_*e*_ and *ν*_*i*_) fluctuates around the stationary levels (see Fig. 3). The Markovian description predicts this phenomena as it contains the impact of finite size effects (the network comprises 10000 neurons). In Fig. 4A, we can see that the distributions of the excitatory and inhibitory population activities are rather well predicted by the formalism: it slightly overestimates the means of the population activities, but it reproduces well the difference of firing between RS and FS cells in the network activity.

**Figure 3:**
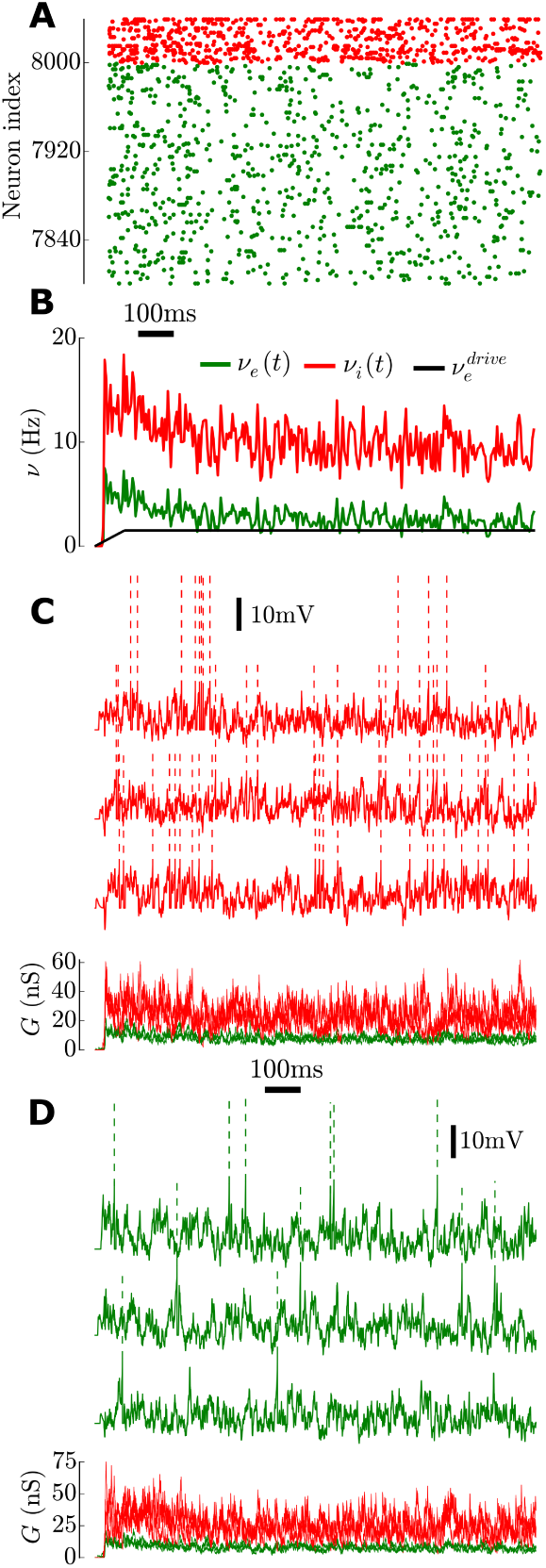
Numerical simulations of the dynamics of a recurrent network of 10000 neurons (see parameters in Table 1) Note that all plots have the same x-axis: time. **(A)** Sample of the spiking activity of 500 neurons (green, 400 excitatory and red, 100 inhibitory). **(B)** Population activity (i.e. spiking activity sampled in 5ms time bins across the population) of the excitatory (green) and inhibitory (red) sub-populations. We also show the applied external drive (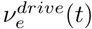, black line), note the slow linear increase to reach 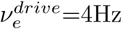 in order to reduce the initial synchronization that would result from an abrupt onset. **(C)** Membrane potential (top) and conductances (bottom, excitatory in green and inhibitory in red) time courses of three randomly chosen inhibitory neurons. **(D)** Membrane potential and conductances time courses of three randomly chosen excitatory neurons.

**Figure 4:**
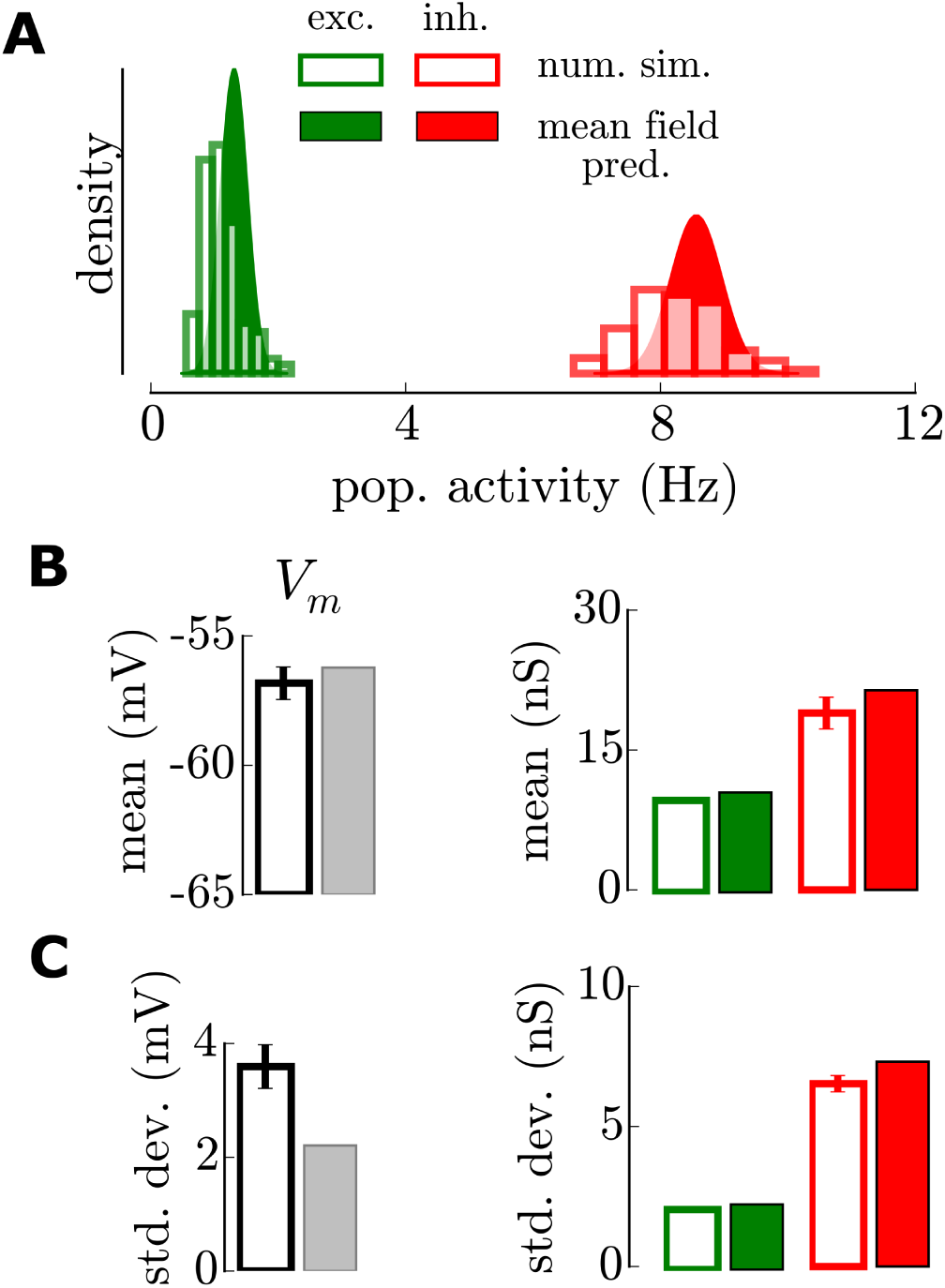
Mean field prediction of the stationary activity. Those quantities are evaluated after discarding the initial 500ms transient. **(A)** Gaussian predictions of the population activities (filled curve) compared to those observed in numerical simulations (empty bars). **(B)** Mean of the membrane potential and conductances time courses. Evaluated over 3 cells for the numerical simulations (empty bars, mean and standard deviation). **(C)** Standard deviation of membrane potential and conductances time courses.

We also investigated whether the average neuronal and synaptic quantities were well predicted by the Markovian formalism. Indeed, we found a very good match for all quantities (see Fig. 4B,C, mean and variance of membrane potential and synaptic conductances). Only the standard deviation of the membrane potential fluctuations was underestimated (Fig. 4C). This discrepancy does not appear detrimental to the formalism as the *V*_*m*_ standard deviation is a key quantity of the transfer function and the formalism still shows a good match. Indeed, this discrepancy might only be due to the presence of threshold-and-reset mechanism or to the low amount of residual synchrony in such finite networks.

### 3.5 Description of the response to time-varying input

We now examine whether the formalism captures the response to time-varying input. Here again, we set the input and examine the response after 500ms of initial simulation to discard transient effects.

We first choose an afferent input of relatively low frequency content (~ [5-20]Hz, *τ*_1_=60ms and *τ*_2_=100ms in Equation 21). The afferent input waveform, formulated in terms of firing rate, was translated into individual afferent spikes targeting the excitatory population. The response of the network to this input is shown in Fig. 5 in comparison with the prediction of the Markovian formalism. The excitatory population activity raises and immediately entrains an increase of the inhibitory population activity. The analytical description captures well the order of magnitude of the deflection, it only slightly underestimates the peak value (Fig. 5B). But the numerical simulations also show a marked hyperpolarization after the stimulation, the return to the baseline level happens only ~ 200-300 ms after the end of the stimulus, and not immediately as predicted by the Markovian framework. Here this strong hyperpolarization is the result of the strong spike-frequency adaptation current in excitatory cells that persists as a repercussion of the high activity evoked by the stimulus. In the Markovian there is no memory of the previous activity and therefore this phenomena can not be accounted for. This typically illustrates a limitation of the analytical description provided here.

**Figure 5:**
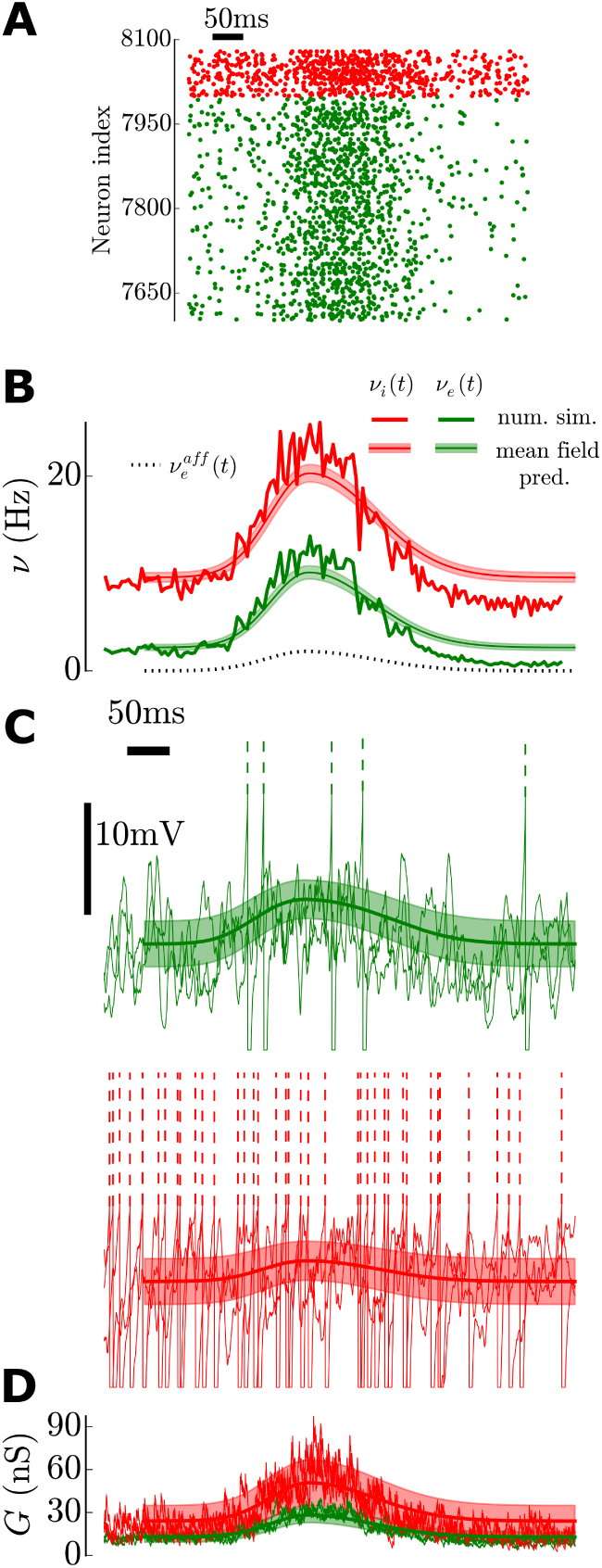
Network response to a time-varying input and associated prediction of the Markovian formalism. For all plots, the x-axis corresponds to time. Shown after 500ms of initial stimulation. **(A)** Sample of the spiking activity of 500 neurons (green, 400 excitatory and red, 100 inhibitory). **(B)** Population activity (in 5ms bins) of the excitatory (green) and inhibitory (red) sub-populations. Superimposed is the mean and standard deviation over time predicted by the Markovian formalism. We also show the applied external stimulation (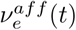, dotted line). **(C)** Membrane potential time courses of three excitatory cells (green, top) and three inhibitory cells (red, bottom) with the prediction of the mean and standard deviation in time. **(D)** Conductance time courses of the six cells in **C** with the predictions of the fluctuations superimposed.

To study more precisely the temporal validity of the formalism, we modulated the network activity by sinusoidal input and compared the response predicted by the analytical description. First, the hyperpolarization phenomena discussed above has a correlate in terms of frequency-dependency. Network activity is overestimated by the mean-field prediction and one can see a discrepancy with respect to numerical simulations at very low frequencies (visible in the [0.01,1]Hz range, see inset in Fig. 6A). Additionally, the numerical simulations showed a marked resonance at ~50Hz. Given the relatively high strength (compared to the external input) of the excitatory-inhibitory loop, the network is close to a bifurcation toward oscillations that are typically in the gamma range (Brunel and Wang, 2003). A sinusoidal input therefore amplifies those frequencies (Ledoux and Brunel, 2011). Because the individual excitatory and inhibitory post-synaptic currents approximately match each other, the theoretical study of Brunel and Wang (2003) would predict oscillations at 50-60Hz (the bifurcation would be achieved by reducing *τ*_*e*_), thus compatible with the present observation. An important insight of this analysis is to show that the network can track very fast temporal variations in the input, even at time scales smaller than the integration time constant of the single neurons (van Vreeswijk and Sompolinsky, 1996). Recurrent neural networks globally behave as low-pass filters (though see Ledoux and Brunel (2011) for a detailed treatment of the appearance of resonances), but with a high cutoff frequency compared to the frequency content of thalamic input for classical artificial stimuli (e.g. in the visual system: drifting gratings, supra-10ms flashes, etc…). Again, *in vivo* experiments in awake mice suggested that V1 cortical networks had a high cut-off frequency (∼100Hz in Reinhold et al. (2015)).

**Figure 6:**
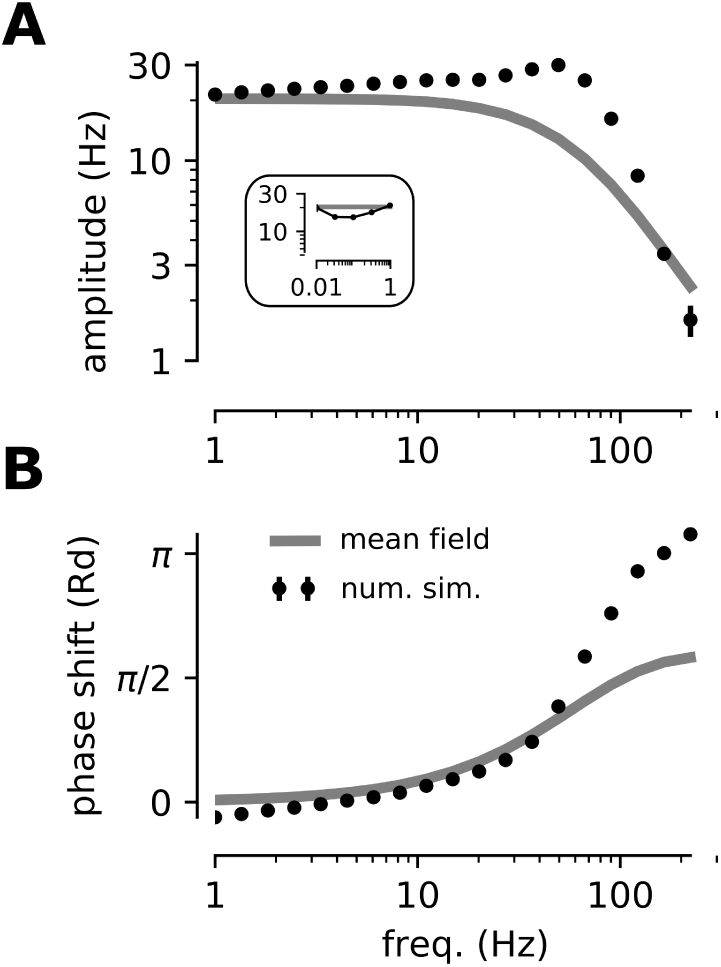
Limitations of the Markovian description in the frequency domain. Response of the network (numerical simulation and analytical description) to sinusoidal stimulation of the form 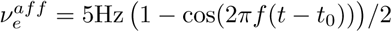. The stimulation was set on at *t*_0_=500ms. The response was fitted by a function of the form *ν*(*t*) = *A* (1 − cos(2*πf*(*t − t*_0_) − *ϕ*))/2. **(A)** Amplitude of the sinusoidal response (*A* in the fitted response) for various frequencies. In the inset, we show the [0.01, 1]Hz range. **(B)** Phase shift of the sinusoidal response (*ϕ* in the fitted response) for various frequencies.

Thus, by comparing numerical simulations of network dynamics and the Markovian formalism, we highlighted the accuracies and discrepancies of this analytical framework to describe both the spontaneous activity and the response of a sparsely connected recurrent network of distinct excitatory and inhibitory cells. We conclude that, given the frequency content of visually evoked network responses in V1 (Muller et al., 2014) (5-20Hz), those limitations would seem to poorly affect the description of such phenomena.

### 3.6 One-dimensional ring model to model VSD imaging

We now embed this local population dynamics description into a spatial model to investigate the emergence of spatio-temporal patterns of activity. The ring model (see e.g. (Hansel and Sompolinsky, 1996)) offers a simple framework to implement such interactions. The local balanced network units are interconnected to each other via two Gaussian connectivity profiles (see Fig. 7 and Methods) according to anatomical connectivity estimates (Buzás et al., 2006). Importantly, we integrate distance-dependent propagation delays due to the finite velocity of axonal conduction of action potentials (see Methods), we took here an axonal conduction velocity of 0.3m/s. Previous theoretical work investigated how the topology of such networks may affect the "macroscopic" quantities of the spiking activity (ensemble correlations and mean firing rates). It was shown that, provided the excitatory-inhibitory balance is the same, those macroscopic properties were globally invariant with respect to the different connectivity patterns (Yger et al., 2011). As the global balance of the ring model is identical to that of the local model (see Methods), those results imply that the *mean-field* analysis of the macroscopic quantities performed in the previous sections provides a good approximation for the dynamics of the topological network considered here. We therefore study the dynamics of the ring model through its *mean-field* description (Equation 23).

**Figure 7:**
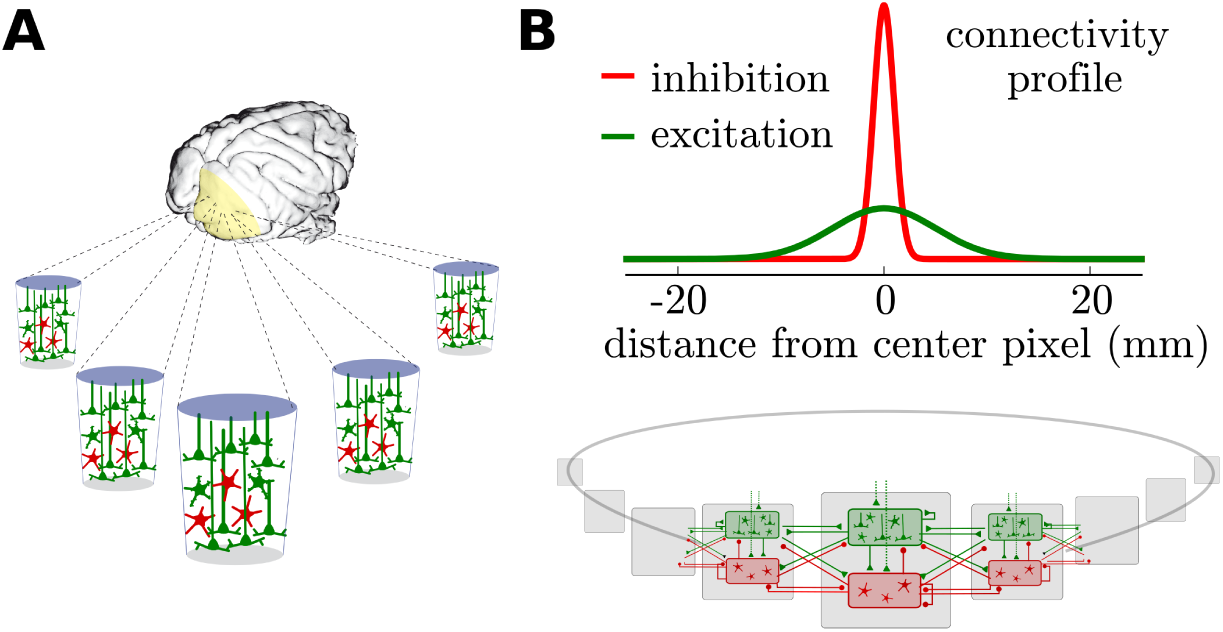
Modeling mesoscopic cortical dynamics. A mesoscopic model of the spatial organization of neocortical populations **(A)** is constructed by inter-connecting the local networks with continuous connectivity profiles of excitatory and inhibitory interactions **(B)**. The lateral connectivity follows two Gaussian profiles of extent *l*_*exc*_=5mm and *l*_*inh*_=1mm for the excitation and inhibition respectively.

We stimulated this large-scale model with an external input mimicking thalamic stimulation. We took a separable spatio-temporal waveform as an input. In space, the profile was a Gaussian curve of extent *l*_*stim*_, in time, it was a piecewise double Gaussian function. This corresponds to the following input:

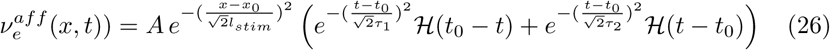

Despite its various amplitude over space (its attenuation from the local maximum), it should be emphasized that this input does not propagate: its maximum is achieved at all position at the same time. To highlight this feature, we implemented a simple analysis of propagation: we normalize the responses with respect to their local amplitude and we look for a specific crossing of the normalized amplitude. To focus on early responses, we highlight the first crossing of the level corresponding to 20% of the maximum amplitude, we will refer to this quantity as the *early response line* (drawn with a white dashed-line, see Fig. 8). In Fig. 8A(i), the horizontal *early response line* indeed shows that the input does not propagate, the fourth of the maximum of the normalized response is achieved everywhere at the same time.

**Figure 8:**
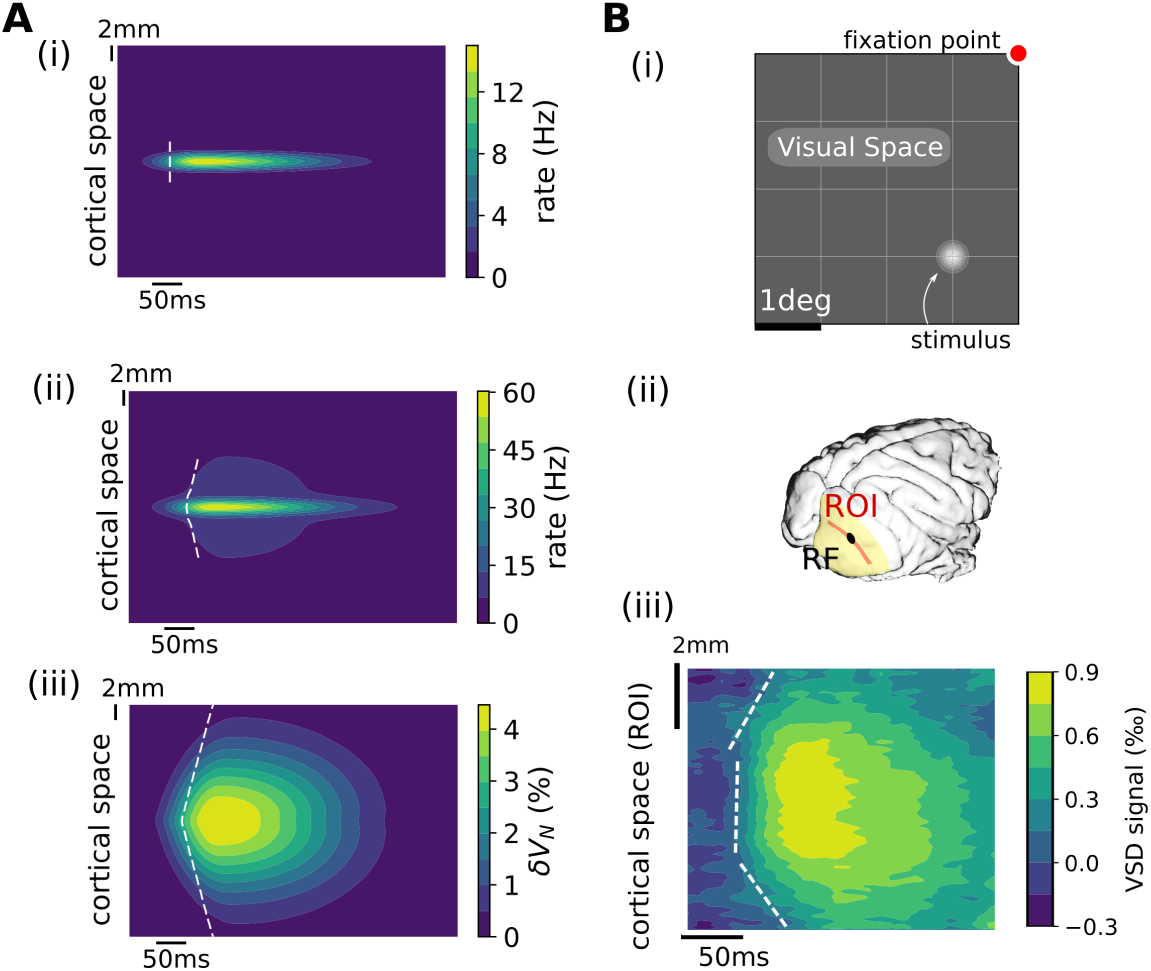
Comparison of evoked activity patterns (A) in the model as a response to an input waveform and (B) observed in awake monkey under voltage-sensitive dye experiments as a response to a visual stimulus. **(A)** (i) Afferent stimulation: an input of the form Equation 26 with the parameters *A*=15Hz, *τ*_1_=50ms, *τ*_2_=150ms and *l*_*exc*_=0.8mm. An *early response line* (white dashed line, see main text) indicates whether the signal exhibits propagation over space (vertical meaning no propagation). Model response in terms of population activity (ii) and (iii) normalized membrane potential (see Methods). **(B)** (i) A gaussian of luminance with angular extent 0.125° is presented in the visual space at 1° (left) and 2° (bottom) from the fixation point. (ii) A one-dimensional region of interest (ROI) is selected surrounding the cortical receptive-field (RF). (iii) VDS imaging response following the visual stimulation. To illustrate the propagation around the center of the evoked response, we arbitrarily splitted the space in three regions (bottom-center-top) and performed a linear fitting over space of the temporal crossing of the 25% level of the local maximum. Note the difference in spatial scale between the model and experiments, the model has here a lower scale to show the spread over the entire ring model.

The response of the model in terms of population dynamics showed a marked propagation (see the V-shape of the *early response line* in Fig. 8A(ii)). This is naturally the result of the local connectivity profiles implemented in the model (see Fig. 1 and Table 1), the excitation has a broad spatial extent, it can depolarizes neighboring locations and evoke spiking (both of excitatory and inhibitory populations). This propagated activity nonetheless exhibits a very strong attenuation over space, this is due to the strong non-linear relationship between depolarizations and firing response. Confirming this picture, the normalized membrane potential responses indeed exhibits the same propagation profile but with a much weaker attenuation over space. Naturally, the propagation dynamics in the model is led by the conduction velocity, see its representation (white dotted line) in Fig. 8A(ii,iii). As expected (Bringuier et al., 1999), the model predicts that the detectability of responses in multiunit recordings have a lower spatial extent than for VSDi responses (see the lower range of the *early response line* that stops when the maximum local response is below 1% of the maximum response).

Fig. 8 also compares visually-evoked propagating waves between the model and VSDi experimnents in the primary visual cortex of awake monkey. A recent phase-based analysis applied at single-trial level (Muller et al., 2014) showed that such propagating waves appear either in spontaneous activity or following visual stimulation. Using a 2-dim space-time representation applied similarly in the data(Fig. 8B(iii)) and in the model (Fig. 8A(iii)) shows that the ring model can reproduce the qualitative features of the propagating wave.

### 3.7 Inferring model parameters from VSDi recordings

In Fig. 9A, we show how the input parameters (the spatial and temporal parameters of the afferent drive: *l*_*stim*_, *τ*_1_ and *τ*_2_) and architecture parameters of the model (lateral propagation speed *v*_*c*_ and extent of lateral connectivity *l*_*exc*_ and *l*_*inh*_) shape the response profile in terms of mean *V*_*m*_ depolarization. Given the previously described qualitative similarity between model and experiments in terms of evoked spatio-temporal patterns, we try to infer the model parameters from the VSDi recordings. To this purpose, we developed an optimization procedure based on a least-square criteria over the spatio-temporal patterns of evoked activity. We examples of the various spatio-temporal patterns recorded in different sessions of VSDi imaging (Fig. 9B, top), we also show how the model was aligned to VDSi recordings and normalized for quantitative comparison of the patterns of evoked activity (Fig. 9B, bottom).

**Figure 9:**
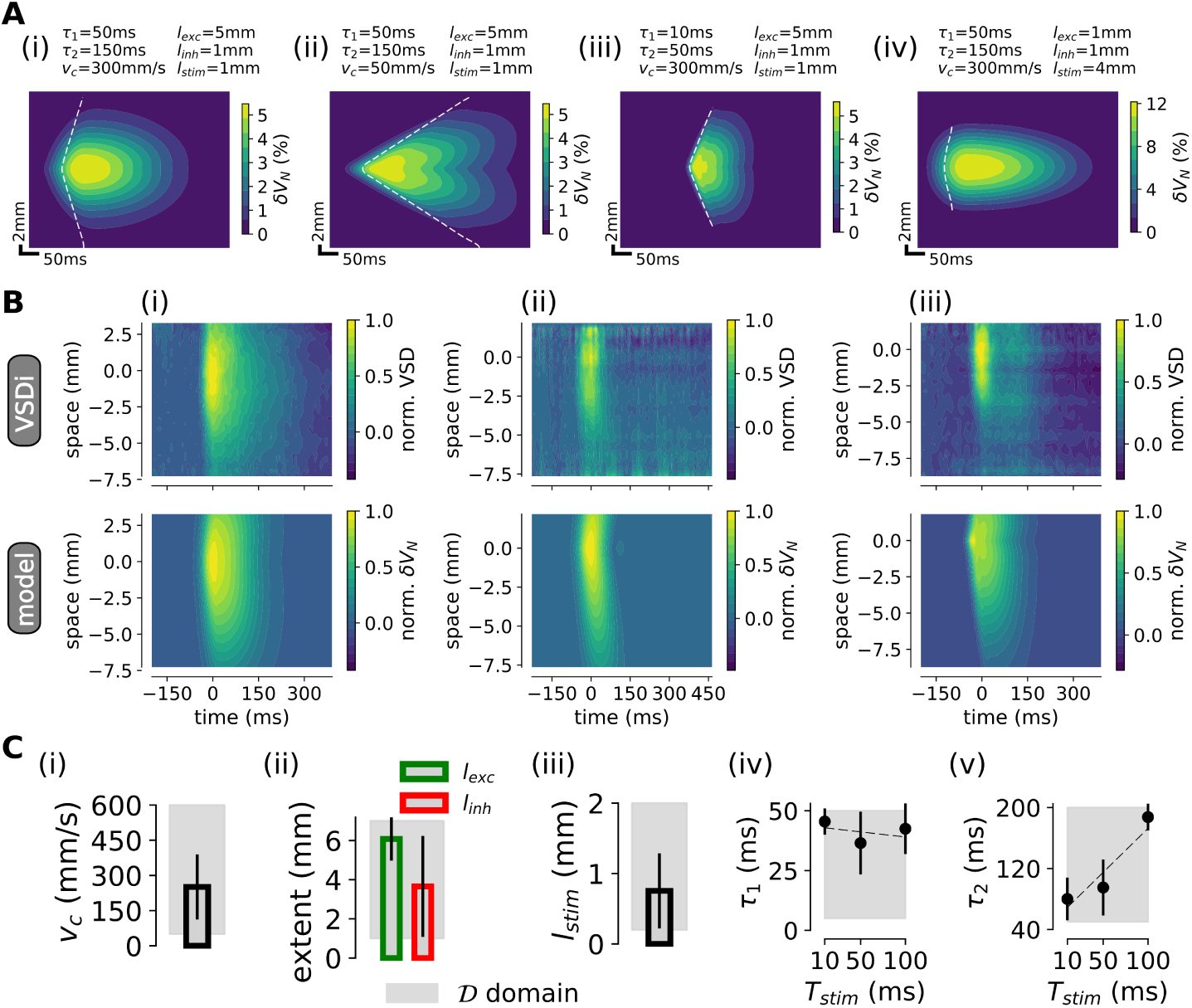
Inferring the model parameters from VSDi recordings. **(A)** Mean depolarization *δV*_*N*_ for different parameters of the model. See parameters values on the figures. From (i) to (ii) we decreased the conduction velocity *v*_*c*_, from (i) to (iii) we decreased the rise *τ*_1_ and decay *τ*_2_ time constants of the stimulus, from (i) to (iv) we enlarged the stimulus extent *l*_*stim*_ and decreased the excitatory lateral conectivity *l*_*exc*_. **(B)** Comparison between data (top) and model (bottom) for the parameters filling the least-square criteria. Shown for three representative recording sessions. Note the common normalization and spatio-temporal sampling that allow to compute the difference between model and experiments (see Methods). **(C)** Estimated model parameters over the whole dataset (n=6 sessions in monkey WA, n=6 sessions in monkey BR): (i) lateral propagation speed *v*_*c*_, (ii) spatial extent of the excitatory (green, *l*_*exc*_) and inhibitory (green, *l*_*exc*_) connectivity profiles, (iii) spatial extent of the stimulus *l*_*stim*_, (iv) onset time constant of the stimulus *τ*_1_ and (v) decay time constant of the stimulus *τ*_2_. We show the domain 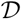 over which the optimization was performed (see Methods).

Overall, we found that the best parameters of the model pointed toward physiologically-realistic values, see Fig. 9C. The optimal speed of lateral propagation was found to be 251 ± 139 mm/s, i.e. close to the ~ 300mm/s value measured experimentally (Waxman and Bennett, 1972). Interestingly, the model predicted one of the key feature of neocortical architecture, the higher lateral extent of the excitatory connectivity (Buzás et al., 2006): we indeed found a significant assymetry between the excitatory and inhibitory extent of lateral connectivity (n=13, p=2e-3, two-sided t-test). Note however that the optimal connectivity values, *l*_*exc*_ = 6.1 ± 1.1mm/s and *l*_*inh*_ = 3.7 ± 2.6mm/s systematically overestimated physiological estimates *l*_*exc*_ = 5mm/s and *l*_*inh*_ ~ 1mm/s in Buzás et al. (2006). The stimulus extent onto the cortical network was found to be 0.8 ± 0.5 mm here underestimating the typical "center response area" (∼2mm, see Fig. 8Aiii). Those two last discrepancies (overestimating lateral connectivity while underestimating stimulus spatial extent) can be explained by the fact that disambiguating between a narrow stimulus with wide lateral propagations and a wide stimulus of narrow lateral propagation is not straightforward (see difference between (i) and (iv) in Fig. 9A). The onset dynamics (captured by the constant *τ*_1_), was found invariant with respect to the length of stimulus presentation (c=0.1, p>0.1, pearson correlation). On the other hand, the decay dynamics was found correlated with the duration of stimulus presentation (c=0.8, p=2e-3, pearson correlation), consistent with the more sustained thalamic activations for longer stimuli presentations as a consequence of the relative linearity of thalamic responses with respect to visual stimulus features (Gawne et al., 1991).

## 4 Discussion

In the present study, we investigated a mean-field model of networks with different electrophysiological properties, described using the AdEx model with conductance-based synapses. We found that the Markovian formalism proposed in El Boustani and Destexhe (2009) was able to describe the steady-state and temporal dynamics of such networks. Though this formalism was shown to be a relatively accurate description of the response simulated in numerical networks, we also showed the limits of this formalism. The relative complexity of the theoretical problem should be stressed: our model includes non-linear phenomena such as spike-frequency adaptation or a voltage-dependent activation curve for spike emission. The proposed semi-analytical approach thus offers a convenient description for theoretical models where an exact analytical treatment would not be achievable.

Unlike previous studies (Brunel, 2000; Vogels and Abbott, 2005; Kumar et al., 2008; El Boustani and Destexhe, 2009), we considered networks of non-linear integrate-and-fire neurons with asymmetric electrophysiological properties between excitatory and inhibitory cells. This type of network is more realistic because it includes the adaptation properties of excitatory cells, and the fact that inhibitory cells are more excitable and fire at higher rates. We could demonstrate the relative accuracy of the Markovian formalism (with the semi-analytical approach) in a situation including this increased complexity. The mean-field model obtained was able to predict the level of spontaneous activity of the network, as well as its response to external time-varying inputs.

This versatile theoretical description of the local cortical network could be improved. For example the strong hyperpolarization of population activity after a transient rise (see Fig. 5B) was shown to be missed by the mean-field formalism. Indeed, this version does not have a memory of the previous activity levels and thus can not account for the effect of the long-lasting spike-frequency adaptation mechanism that has been strongly activated by the activity evoked by the stimulus. One could design another version of the Markovian formalism to capture such adaptation-mediated effects. Instead of accounting for adaptation within the *transfer function* (i.e. accounting only for its stationary effects), one can introduce a new variable with a dependency on time and activity: a “population adaptation current”, that can directly be derived from the equation of the AdExp model. Additionally, recent semi-analytical work (Augustin et al., 2016) in current-based networks yielded very accurate descriptions of network activity both at low and high frequency content, translating those results to conductance-based networks could overcome the limitations of our description. Investigating such formalisms and their accuracy should be the focus of future work.

We further showed that the present mean-field approach can be used to model VSDi data. Not only the present mean-field framework gives access to the mean voltage and its time evolution, but it can easily be extended to model VSDi signals. The present model represents a local population of cortical excitatory and inhibitory neurons, and thus can be thought to represent a “pixel” of the VSDi. The full VSDi model was obtained by embedding the present local population description within a spatial model, under the form of a ring-like arrangement of RS-FS mean-field units (see Fig. 8). In this simple model, a localized input led to propagating-wave activity, very similar to experiments (see Fig. 8). This demonstrates that the present mean-field approach can be used to model VSDi experiments. This study thus constitutes a "proof of concept" validated on the spatio-temporal pattern of neocortical activity evoked by a single stimulus. Investigating whether the present theoretical model yield deeper insight into neocortical computation is the focus of current work.

## Acknowledgments

Research supported by the CNRS, the ICODE excellence network, the European Community (Human Brain Project H2020-720270 to A.D., FET Grant BrainScaleS FP7-269921 to A.D. and F.C) and the ANR BalaV1 and Trajectory (ANR-13-BSV4-0014-02 to F.C). Y.Z. was supported by fellowships from the Initiative d’Excellence Paris-Saclay and the Fondation pour la Recherche Médicale (FDT 20150532751).

